# Disrupted microglial iron homeostasis in progressive multiple sclerosis

**DOI:** 10.1101/2021.05.09.443127

**Authors:** Jonathan D. Proto, Mindy Zhang, Sean Ryan, Xinting Yao, Yinyin Huang, Yi-Chien Chang, Michael R. Dufault, Emilie Christie, Anthony Chomyka, Jackie Saleh, Jose Sancho, Timothy Hammond, Bruce Trapp, Dimitry Ofengeim

## Abstract

Multiple Sclerosis (MS) is a chronic autoimmune disease affecting the central nervous system (CNS). Despite therapies that reduce relapses, many patients eventually develop secondary progressive MS (SPMS), characterized by ongoing and irreversible neurodegeneration and worsening clinical symptoms. Microglia are the resident innate immune cells of the CNS. While the cellular and molecular determinants of disability progression in MS remain incompletely understood, they are thought to include non-resolving microglial activation and chronic oxidative injury. In this study, our aim was to better characterize microglia in SPMS tissues to identify disease-related changes at the single cell level. We performed single nucleus RNA-seq (snRNA-seq) on cryopreserved post-mortem brain cortex and identified disease associated changes in multiple cell types and in particular distinct SPMS-enriched microglia subsets. When compared to the cluster most enriched in healthy controls (i.e. homeostatic microglia), we found a number of SPMS-enriched clusters with transcriptional profiles reflecting increased oxidative stress and perturbed iron homeostasis. Using histology and RNA-scope, we confirmed the presence of iron accumulating, ferritin-light chain (*FTL*)-expressing microglia *in situ*. Among disease-enriched clusters, we found evidence for divergent responses to iron accumulation and identified the antioxidant enzyme GPX4 as a key fate determinant. These data help elucidate processes that occur in progressive MS brains, and highlight novel nodes for therapeutic intervention.

## INTRODUCTION

Multiple sclerosis (MS) is a neuroinflammatory disease that leads to progressive white matter and gray matter loss^1^. About 85% of MS patients are initially diagnosed with a relapsing-remitting form of disease (RRMS). Despite the availability of disease-modifying treatments (DMT) that reduce relapses, many patients eventually develop secondary progressive MS (SPMS), characterized by ongoing neurodegeneration resulting in irreversible neurological disability. The cellular/molecular determinants of disability progression remain incompletely understood. At disease onset, demyelination results from immune-mediated cytotoxic damage to oligodendrocytes (OLs) in the CNS. Indeed, the success of DMTs targeting peripheral immunity illustrates the causative role of these populations in disease pathogenesis. Accumulating evidence suggests that the involvement of innate immune cells in this process drives progressive axonal damage in the CNS white matter^2^. Studies have identified a rim of microglia along the lesion border that is believed to participate in deleterious processes in the CNS. While the presence of innate immune cells in the lesion border, but the full identity and function of these cells has remained elusive^3^.

Advances in single cell analytical approaches in the CNS have allowed us to better understand various aspects of neuro-immune interaction leading to progressive neurodegeneration^4^. Single nucleus RNA sequencing (snRNA-seq) is a powerful approach to analyze samples not amenable to cell sorting, such as cryopreserved brain tissue. Several studies have examined this in the animal models of MS as well as in patient derived samples^5–8^. Much of the focus has been the role of a particular lineage in the context of MS, such as oligodendrocytes or neurons^7,8^. In this study we employed a modified version of a previously described snRNA-seq and paired it ^9,10^ with bulk RNA sequencing to interrogate cellular changes in progressive MS. We were able to definitively resolve brain-resident populations (neurons, astrocytes, OLs, microglia) and identify several disease-associated phenotypes. While innate immune cells represented only a small fraction of cells identified by snRNA-seq, genes associated with microglia account for the majority of transcriptional changes in bulk tissue RNA-seq analysis. Based on our data, as well as published literature, a key driver of disease progression in MS is an alteration in the neuroinflammatory milieu in the CNS; consistent with this we identified a robust microglial-specific signature in the MS bulk RNA-seq analysis. To better understand the role of these cells in the CNS during disease progression, we utilized a single-nuclei approach from post-mortem brain tissue. We used an iterative analytical approach to identify disease associated transcriptional changes in microglial cells across several batches of high-quality samples and were able to identify microglial populations associated with MS, which we describe as MS-MG1 and MS-MG2. Using these observations, we show that these two populations are spatially distinct in SPMS white matter lesions. We further suggest that a microglial response to iron may be a key driver of some of the disease associated phenotype. Consistent with previous data, we show accumulation of ferritin light-chain gene expression in microglia in white matter lesions^11^. Using, our snRNA-seq data set, we can identify a population of microglia that are responding to iron, and thereby characterize their transcriptomic profile. Furthermore, we show that perturbing iron handling in microglia *in vitro* can lead to an altered cellular phenotype. These findings not only underscore the importance of utilizing single cell approaches to better understand disease pathology in MS but point to an important role for iron handling by microglial cells in the CNS in MS as well as other neurodegenerative diseases.

## RESULTS

### Major CNS cell types were identified in brain white matter from SPMS and healthy donors

Nuclei were isolated from high quality samples (RNA integrity number (RIN) of >6.5) of lesioned or normal appearing white matter (NAWM) derived from secondary progressive MS patients (n=7) or neurological controls (n=8) **(Supplementary table 1**). Staining for NeuN, a neuron-restricted nuclear membrane protein^9,10^, confirmed that samples were enriched for non-neuronal nuclei, as expected for white matter derived cells **(Figure 1A**). Pilot experiments confirmed that our procedure maintains the RNA integrity and transcriptional fidelity as compared to bulk tissue analysis (**Figure S1**). Using our optimized isolation method, we were able to detect >500 genes across the various cell types (**Figure S2**). After rigorous pre-processing and quality control, we obtained a total of 124,934 single cell nuclear transcriptomes. To identify major CNS cell types, including neurons, astrocytes, microglia, oligodendrocytes (OL), and oligodendrocyte progenitor cells (OPCs), nuclei were scored for enrichment of cell-type specific transcriptome profiles, which consisted of 13-21 marker genes per cell type (**Supplementary table 2)**. These major cell types were identified in nearly all patients, as well as cells scoring high for both OL and OPC markers, which we subsequently refer to as OL-OPC **(Figure 1D**). Of note, similar numbers of each cell type were derived from control or SPMS patients. In previous studies, this has been a particular challenge in regards to microglia ^12,13^**(Figure 1E).** In comparison to other CNS cell types, we found significantly fewer counts in microglia, a finding consistent with previous studies (**Figure S2B).** Others have reported that well characterized human microglial activation genes (*CD74*, *SPP1*, *TMSB4X*, *CST3*, *APOE*, *HSPA1A*) are depleted in nuclei, but we did not find a decrease in read counts when we compared bulk tissue to nuclei RNA-seq (**Figure S1**).^14^

**Figure 1.**
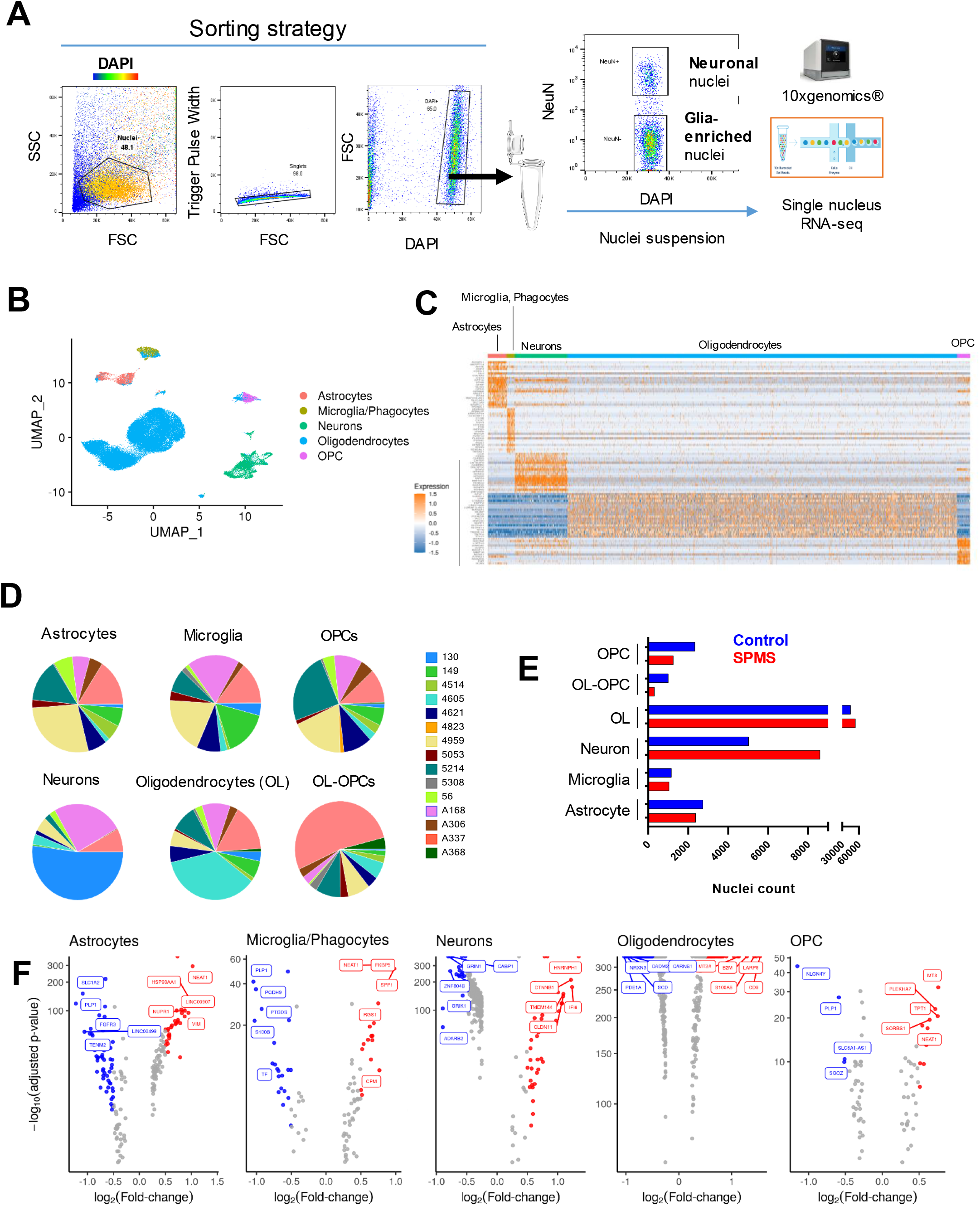
snRNAseq approach. (**A**) Schematic of nuclei sorting strategy. (**B)** UMAP visualization of major CNS cells types and (**C**) heat map of gene expression across annotated cell types. (**D**) Pie charts of patient contribution per cell type and (**E**) distribution by disease status (n=7 SPMS, 8 neurologic controls). (**F**) Volcano plot of DEG across each cell type by disease status (MS vs. Healthy).

### Common features of microglia subpopulations in progressive MS patients

We next examined disease-associated changes across cell types. In SPMS patients, both OL and astrocytes downregulated genes related to axonogenesis, while genes related to anti-apoptotic processes, ATP synthesis, and cytokine signaling were among those upregulated (**Figure 1F, Figure S2C**). Non-resolving neuroinflammation driven by innate immunity (i.e., microglia/macrophages) has been increasingly implicated in conversion to progressive MS^12,13^. While microglia represent only a small fraction of the cells within the CNS, microglial-enriched genes dominated gene expression changes in bulk sequencing experiments. To identify disease-associated changes in microglial transcriptomes, we first examined a *discovery set* of 421 microglia derived from two control and two MS patients. Using a graph-based clustering approach, we identified three broad microglial clusters (**Figure 2A**). NAWM control-derived microglia were localized to cluster 1 (hereafter referred to as *HC-g1)* and two clusters were highly enriched in SPMS samples (Clusters 2 and 3, hereafter referred to as *MS-g2* and *MS-g3,* respectively). More than 60 genes were differentially expressed in *both* MS-enriched clusters versus healthy, indicating there are common features in SPMS microglia as they shift from a homeostatic phenotype (**Figure 2B**). Of these shared genes, upregulated transcripts included those associated with cytoskeletal organization (*CFL1, ACTB, TMSB10, TMSB4X*) and cellular iron homeostasis (*HAMP, FTL, FTH1*). While some components of immunoglobulin and cytokine/chemokine-mediated signaling were upregulated (*FCGR3A*), others were notably downregulated (*CSF2RA)* (**Figure 2B).** Interestingly, homeostatic genes, including *CX3CR1*, *P2RY12*, *CSF1R*, and *NAV2*, were only downregulated in the MS-g2 cluster (**Figure 2C,D).**

**Figure 2.**
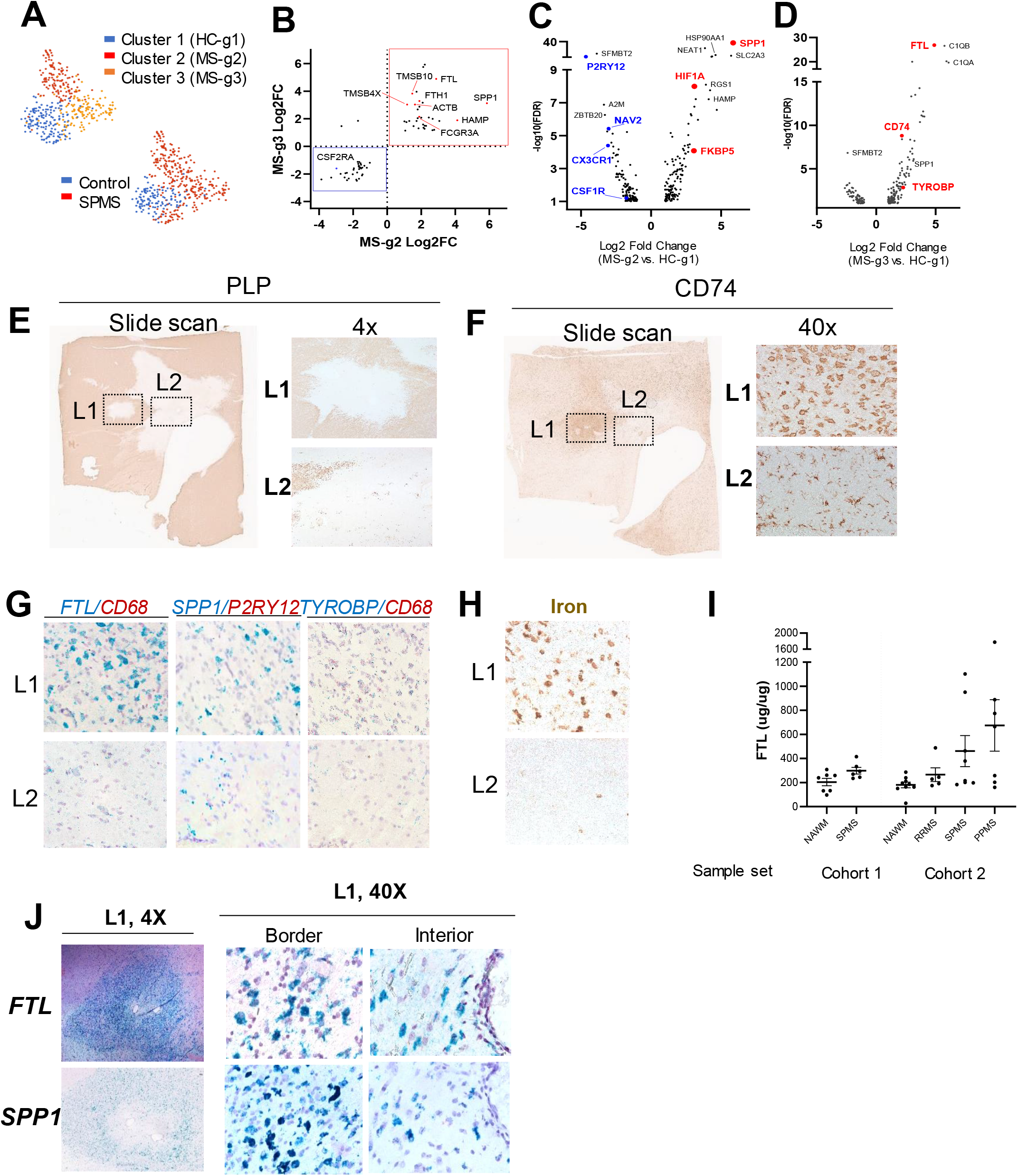
M S-associated subsets identified and validated in situ. (**A**) UMap of discovery set (421 cells). **(B**) Fold-changes in DEG changed in both MS-g2 (x-axis) and MS-g3 (y-axis). **(C)** Volcano plots depicting DEG in MS-g2 and **(D)** MS-g3 versus HC-g1. (**E**) IHC staining for PLP and (**F**) CD74 in a periventricular WM lesion (**left,** slide scan; **right**, 4X). (**G**) *In situ* staining of *FTL, SPP1, and TYROBP* transcripts within lesions (40x). (**H)** Cytoplasmic accumulation of iron visualized by DAB-enhanced Turnbull staining (40x). **(I)** Measurement of FTL in samples from control or diseased tissues (n= 15 control, 25 MS)**. (J)** In situ labeling of *FTL* and *SPP1* both at the lesion border and interior.

### *FTL* and *SPP1* expression observed to be spatially distinct within an active lesion

Genes previously associated with a disease associated phenotypes in MS such as *SPP1*, *CD74,* and *FTL* were among those distinguishing between the MS-g2 and MS-g3 populations (**Figure 2C,D**)^15^. While our data identified at least two major clusters of disease-associated microglia in MS brains, the spatial distribution of these cell types in lesions is unclear. We therefore examined tissue from the temporal lobe subcortical white matter from an SPMS patient containing both an active (L1) and inactive lesion (L2) (**Figure 2E**). CD74+ cells in L1 had an amoeboid morphology, while those in L2 were more ramified (**Figure 2F**). Staining for both *FTL* and *SPP1* was prominent in the active lesion (L1) and sparse in the inactive lesion (L2, **Figure 2G**). Within L1, *FTL* staining was prominent throughout the lesion. In contrast, *SPP1*+ cells increased towards the border of the lesion. (**Figure 2J**). Based on these data, *SPP1* and *FTL* may be enriched in spatially distinct phagocytes within an active lesion.

### Sub-clustering reveals unique disease-enriched microglia populations

Cellular iron, primarily cytoplasmic, was prominent in L1 and not L2 (**Figure 2H**). Interestingly, FTL protein was elevated in progressive MS samples at the tissue level, which may partially reflect lesion load (**Figure 2I**). Furthermore, the role of *FTL* and the microglial response to iron in MS lesions may be a critical determinant of lesion status. We then expanded analysis to our entire dataset of over 2,000 microglia/phagocytes. From these larger dataset, we identified a total of 11 clusters, each containing cells from ≥8 individuals and ranging from 711 to 29 cells each. Control-derived microglia accounted for the majority of 5 clusters (0, 1, 4, 7, 10) and SPMS-derived cells were primarily found in 5 clusters (2, 3, 5, 6, 8) (**Figure 3A**). We also identified CNS macrophages (cluster 9) based on expression of *CD209*, *F13A1*, and *MRC1* ^16^. To identify whether the 10 microglia clusters shared characteristics of cell states identified in the discovery set, we scored each cluster for enrichment of the HC-g1, MS-g2, and MS-g3 profiles (**Figure 3B-D)**. The HC-g1 and MS-g3 signature was especially enriched in clusters 0 and 6, respectively (*hereafter referred to as G1 and G3*). In contrast, the MS-g2 profile was enriched in three clusters (4, 5, and 8; *hereafter referred to as G2-a, −b, −c*) **(Figure 3C)**. Notably, G2 (a, b, c) and G3 microglia were found in 13 and 8 individuals, respectively (**Figure S3B**). Thus, we validated the profiles identified in the discovery set are independent of individual-driven bias and were able to expand the initial definition of the MS-g2 state into three additional groups (G2-a, −b, −c).

**Figure 3.**
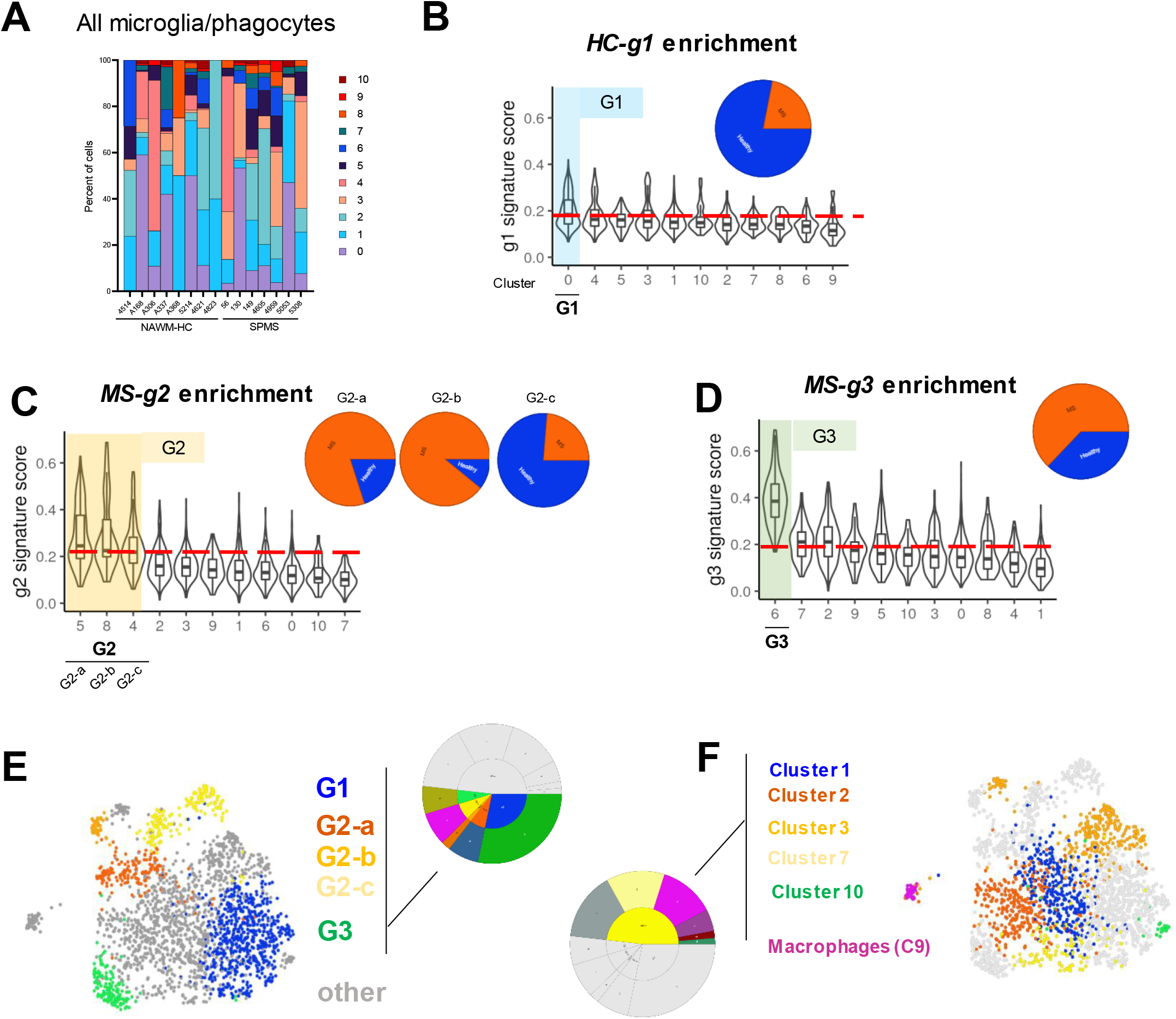
M S-associated subsets identified across patient samples. (**A**) Distribution of clusters by patient. **(B**) Enrichment for discovery-set derived profiles and distribution between control and SPMS samples (inset pie chart) for HC-g1, (**C**) MS-g2, and (**D**) MS-g3. (**E**) UMAP colored by inferred identity from discovery set and (**F**) all unmapped clusters.

G1, G2, and G3 clusters may represent distinct cell fates, with the remaining five clusters falling within this spectrum or standing apart (**Figure 3E,F**). We used multiple approaches to try to better understand the functional role of the different microglial clusters identified in our data set. We first used published mouse data sets to determine the function of each cluster. Transcriptional signatures have been described for various microglia subclusters in murine models of disease, including neurodegenerative or disease-associated microglia (DAM), interferon-responsive (IFN-R), and antigen presenting (MHC-II high, MHCII+) related populations^17^. In contrast to murine models, these profiles did not discriminate as well between microglia from healthy and diseased patients. A number of mouse disease-associated signature genes were highly expressed in G1 (i.e. homeostatic) cells (**Figure S3A, C)**. Likewise, human monocyte-derived macrophages (hMDM) stimulated with LPS (10 ng/mL), an often-used model system for inflammatory-type macrophages, also did not recapitulate the cell states observed in SPMS patients. Rather, relative to homeostatic (G1) microglia, this gene set was most enriched in cluster G2-c. In contrast to G2-a and −b, this cluster was *not* enriched in MS patients (**Figure S3B,D**). This illustrates that a widely used *in vitro* model of inflammatory-type macrophages does not adequately capture the transcriptional responses of microglia in the setting of progressive MS.

### Transcriptomic data and *in vitro* studies link cellular iron, oxidative stress, and survival pathways

Based on gene ontology annotations, G2 and G3 transcriptional states may reflect the interplay of cellular stress, iron metabolism, and inflammation pathways (**Figure 4A)**. Alterations in cellular iron handling have been implicated in progressive MS pathology as well as Alzheimer’s disease^13,18^. Compared to homeostatic microglia, genes encoding components of cellular iron metabolism were enriched in subsets of disease associated cells. Components of ferritin, *FTL* and *FTH1*, were elevated in G3 microglia (**Figure 4B**). In contrast, the transcript for hepcidin (*HAMP*), a secreted molecule that negatively regulates iron uptake, was most elevated in G2-a. Likewise, the iron transporters *SLC11A1* and *SLC25A37* and the heme-clearing enzyme *HMOX1*, were upregulated in G2-a clusters, and not in G3 cells. Localized to late endosome and lysosomes, *SLC11A1* pumps iron from phagosomes into the cytosol. In contrast, SLC25A37 transports iron into the mitochondria where it can be used for heme synthesis.

**Figure 4.**
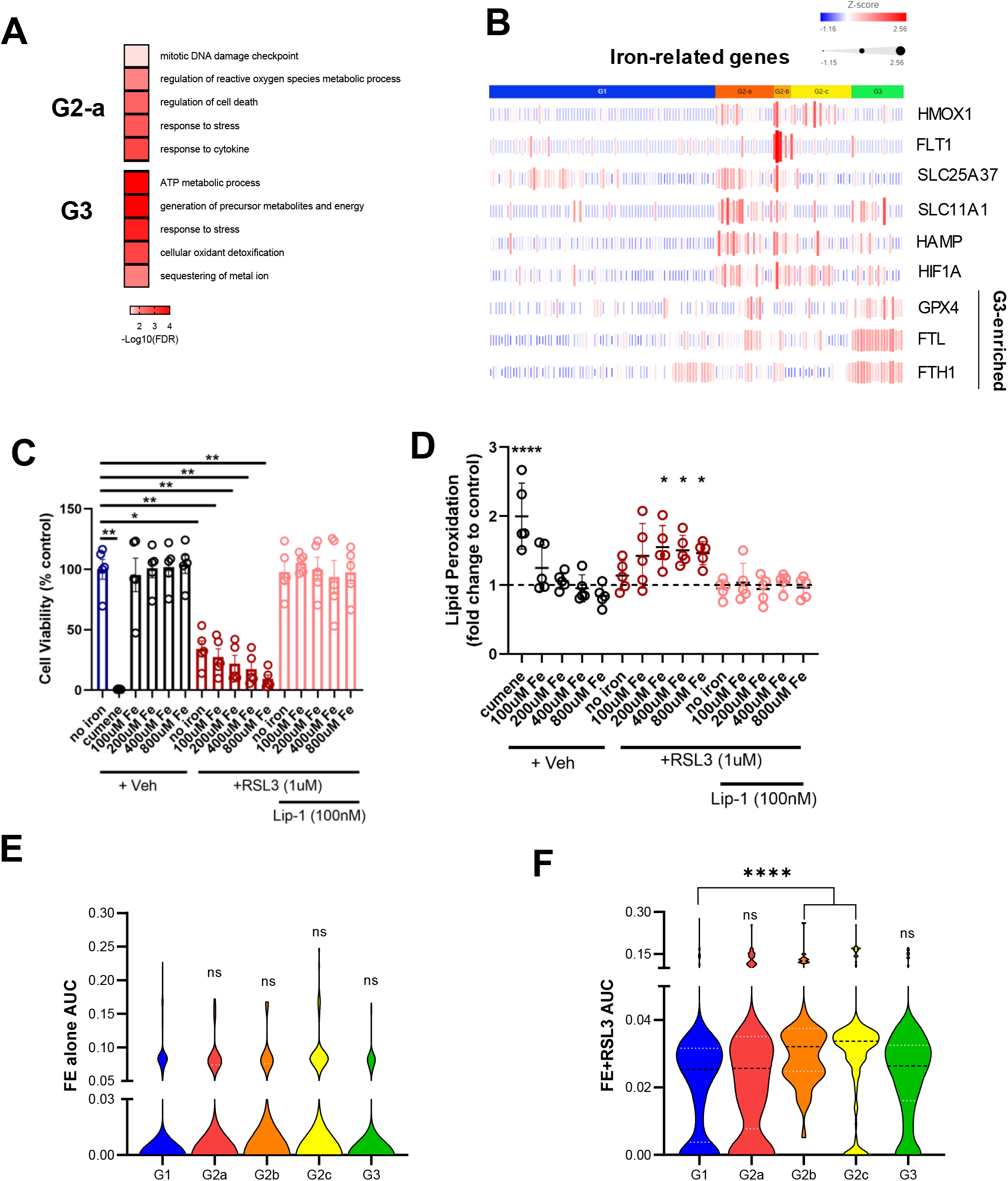
Genes upregulated in MS-MG2 promote survival in a human microglial cell line following iron loading *in vitro*. (**A**) Select gene ontology annotations for DEG from G2-a and G3 clusters. (**B**) Heatmap depicting expression of iron-related genes. (**C**) Cell death following treatment with RSL-3 +/− Liproxstatin 1 in iron loaded cells (n=5 independent experiments, RM one-way ANOVA with Dunnett’s post-hoc analysis). (**D**) Quantification of BODIPY staining for lipid peroxidation (n=5 experiments, Two-way ANOVA with Dunnett’s post-hoc analysis). (**E**) AUCell score distribution for iron-loading signature alone and (**F**) with RSL3 treatment (****p<0.0001, *p<0.05; Kruskall-Wallace ANOVA with Dunn’s post-hoc analysis).

When present in excess amounts, insufficiently sequestered cellular iron results in oxidative stress and cellular damage. Although iron-related pathways were upregulated in both G2 and G3 states, the stress-induced transcription factor *HIF1A* and its target genes (*HMOX1, FLT1*) were markedly upregulated in G2 cells (**Figure 4B**). In contrast, G3 cells demonstrated upregulation glutathione peroxidase 4 (GPX4)^19^. To better understand how this system functions in microglia, we loaded a human microglia cell line with iron in the presence or absence of the GPX4 inhibitor RSL3^20^. Iron-loading alone had no effect on cell viability, but when combined with GPx4 inhibition, there was a significant increase in cell death. This effect could be rescued by treatment with liproxstatin-1 (**Figure 4C**). Staining with BODIPY confirmed lipid peroxidation in this model was dependent on GPX-4 activity. (**Figure 4D**). Using this model system, we examined the transcriptional response to iron-loading with (FE+RSL3) and without (FE alone) GPX4 inhibition. Iron loading alone upregulated 79 genes. However, when GPX4 was inhibited, a total of 156 genes were upregulated (**Figure S4**). We scored all microglia for enrichment of the top 15 DEG. The “FE alone” gene set was not significantly enriched in any clusters, although G2c showed a trend (p=0.053). However, the FE+RSL3 gene set was significantly enriched in all G2-a and G2-b subsets (**Figure 4E,F**). These data show that disrupting antioxidant responses during iron loading *in vitro* results in cell death and suggest that an altered response to iron by microglia may be a driver of oxidative stress and disease progression in MS.

### SPMS-associated microglia gene signatures are detected in tissue samples from other neurodegenerative diseases but not in psychiatric patients

Previous studies show that there is a dominant immune signature in bulk tissue from various neurodegenerative disease^21^. These observations reflect the change in cellular composition in the context of disease, however the ability to identify cellular phenotypic changes within these data sets has not been straightforward. While bulk sequencing experiments have been used extensively to characterize disease mechanism, the ability to deconvolute these types of complex signatures from bulk sequencing experiments has been challenging. We performed bulk sequencing on an independent set of brain tissue sections from SPMS, PPMS, RRMS, and control patients (**Supplementary table 3**). Canonical pathway analysis of differentially expressed genes (DEG) indicated clear alteration of immune function in MS samples (**Figure 5A**). We performed gene set variation analysis (GSVA) to predict relative enrichment of immune populations across all samples. Not surprisingly, we found an enrichment of lymphocyte (B and T cell) and microglia cell signatures in diseased patients and a corollary decrease in the neuronal signature (**Figure 5B**). Thus, changes in the tissue transcriptome likely reflect both infiltration of peripheral cells (i.e. lymphocytes, monocytes) and activation of brain-resident populations, including microglia and CNS macrophages.

**Figure 5.**
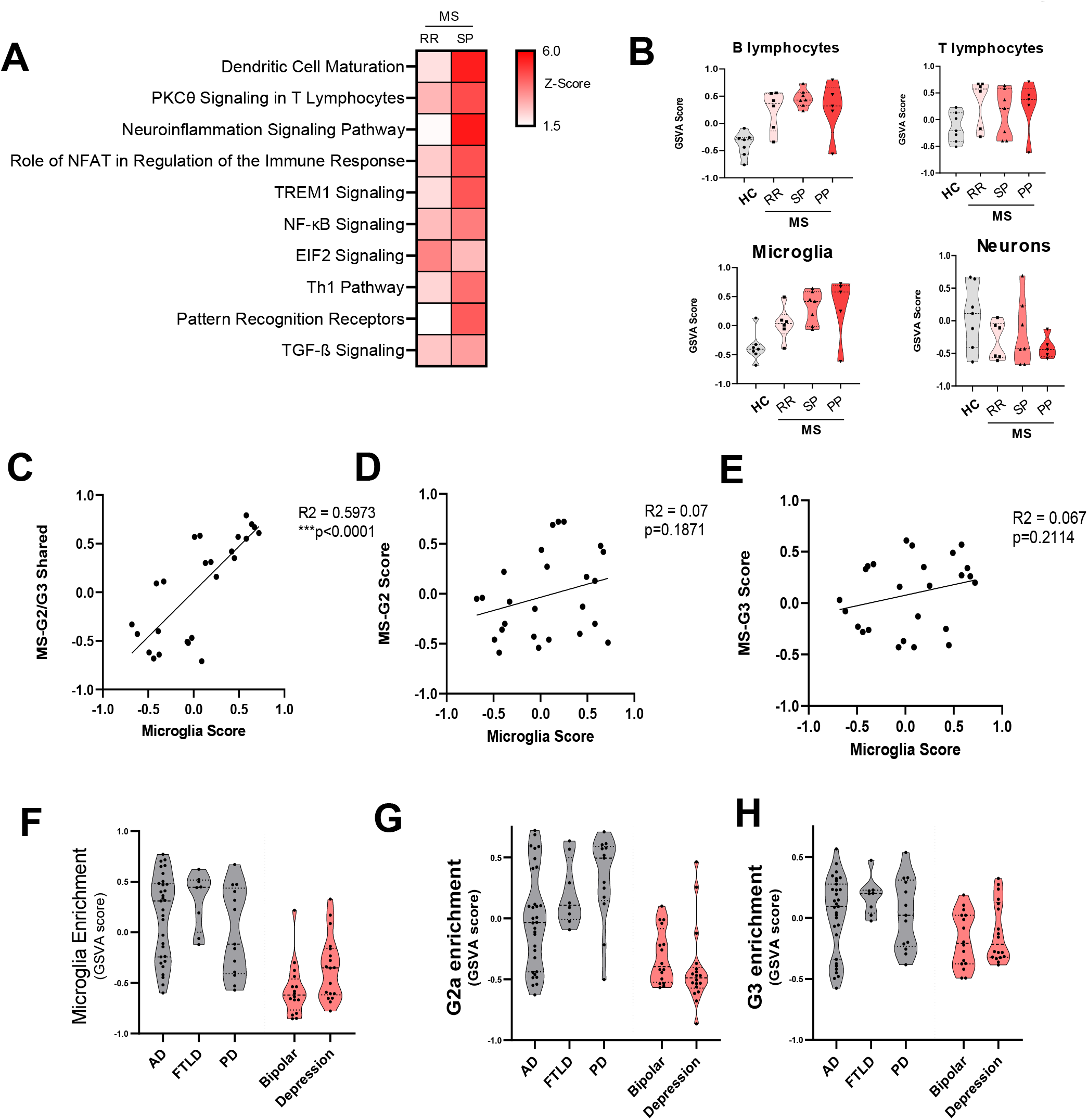
Tissue level transcriptomic changes largely reflect disease-related changes in microglia. (**A**) Heatmap of canonical signaling pathways in SPMS and RRMS samples. (**B**) Enrichment analysis for cell specific signatures. (**D**) Correlation testing between G2/G3 shared, (**E**) G2-a, and (**F**) G3 scores with overall microglia scores. (**F**) Similar scoring across several indications for overall microglial enrichment, (**G**) G2a, and (**H**) G3 signature enrichment. RR, relapsing-remitting MS; SP, secondary progressive MS

We next used a similar approach to look at relative enrichment of the G2-a and G3 microglia states identified in the sNuc-seq data, including the transcriptional changes they have in common (G2/G3 common). Interestingly, individual G2-a and G3 scores did not correlate with the baseline microglia score. In contrast, the G2/G3 common signature was correlated with the microglia score. These results may reflect inter-patient differences in the proportions of G2-a and G3-like microglia (**Figure 5C-E**). We next applied a similar approach using public datasets covering a number of neurodegenerative (Alzheimer’s disease, fronto-temporal dementia, and Parkinson’s disease) and psychiatric indications (bipolar and major depression) (**Figure 5F–H, Supplementary Table 5**. These signatures were present in other neurological diseases as well, particularly neurodegeneration. Although additional studies will be required, this suggests that there are similarities between microglial responses across neurodegenerative diseases and variability among patients may reflect differences in underlying microglia pathology.

## DISCUSSION

While disease driven transcriptional alterations occur in all cell types of the CNS, microglial phenotypic changes may be of particular importance in the context of MS pathology. Innate immune processes both in the periphery and in the CNS of MS patients have emerged as the next wave of targets to alter disease progression. Understanding the role for microglia in both supporting normal CNS homeostasis and mediating deleterious or protective processes in the context of MS is critical in identifying transformational interventions for progressive disease. In this study, we used an unbiased single cell sequencing approach to identify changes in microglial phenotypes in progressive MS, and identified a role for iron and oxidative stress response as a key regulator in determining the microglial phenotype. We employed a rigorous approach to sequence single nuclei from high-quality/rapid-autopsy white matter tissue from secondary progressive MS patients. While previous studies have focused on the role for oxidative damage in mediating axonal and neuronal pathology in MS^22^, our data suggest that changes in iron metabolism leads to oxidative stress that causes and altered inflammatory phenotype in microglial cells. These data suggest that microglia are a key cell type that responds to MS-induced oxidative stress in the CNS, that has previously been described to occur in disease progression^23^.

A finding in our study was that the HIF1A and iron-induced response are coupled in at least a subpopulation of microglial cells in MS. This is consistent with previous findings in other neurodegenerative diseases such as Parkinson’s disease^24^. The interplay of these two nodes in regulating CNS homeostasis needs to be further explored in subsequent studies, and in particular in progressive MS. Interestingly, iron mediated signaling can lead to an inflammatory form of cell death, called ferroptosis and its role in MS should be further explored^25^. This form of cell death is regulated by GPX4, which is a enzyme responsible for blocking the deleterious iron-induced cellular response and is altered in our MS transcriptomic data sets^26,27^. Iron accumulation has been observed in MS as well as in other disease both histologically and by MRI; an intriguing hypothesis is that degeneration in the CNS is coupled to this accumulation, in conjunction with GPX4 dysfunction.

An additional finding from our study is the observation that microglial cells alter their profiles in different ways to respond to disease progression induced damage. While our study clearly identifies multiple microglial subtypes that emerge in the diseased brain, the relationship between each of these clusters and the trajectory by which immune cells in the CNS alter their transcriptional signature to acquire these phenotypes remains to be fully explored. An interesting therapeutic question based on this observation of multiple microglial clusters is the possibility to independently target one subtype of cells to achieve therapeutic benefit. In our study, we employ an AUC approach to characterize certain disease phenotypes and show that we are able to infer their presence in other data sets. In particular, we are able to identify altered microglial states in other neurodegenerative diseases, using available bulk tissue analysis. In subsequent studies it will be important to test these phenotypic predictions by employing single cell analysis.

## METHODS

### Patient samples

Cryopreserved post-mortem brain specimens were obtained from the UCLA Brain Bank or the Cleveland Clinic Rapid Autopsy Program^28^. Tissue specimens consisted primarily of subcortical or periventricular white matter with a smaller portion of the surrounding gray matter. Available clinical data and specimen details are provided in Supplementary Table 1 and Supplementary Table 3 for samples used for snRNA-seq and bulk RNA-seq, respectively. For snRNA-seq, Neurological controls included 5 females and 3 males (65 years old +/− 15.5 years) and SPMS patients included 6 females and 1 male (59 years old +/− 8.2 years). For bulk RNA-seq, performed with a separate set of samples, neurological controls included 3 females and 4 males RRMS patients included 2 females and 4 males, SPMS samples included 5 females and 2 males, and PPMS included 3 females and 2 males.

### Nuclei preparation

Suspensions of nuclei were prepared from homogenized tissue following a previously described method with modifications^10^. In brief, approximately 200-400 mg of tissue was processed on ice in a nuclei isolation buffer consisting of 240 mM sucrose, 24 mM KCL, 4.8 mM MgCl2, 9.59 mM Tris (pH 8.0), 0.97 uM DTT, 0.1% (v/v) Triton X-100, 1x protease inhibitor (Fisher, PI78429), 0.4 units/uL RNAseIn (Fisher, PR-N2511), and 0.2 units/uL Superasin (Fisher, AM2694) in nuclease free water. Samples were first placed in a petri dish containing buffer and minced using a razor. The tissue suspension was then then homogenized using a glass dounce homogenizer and the homogenate was filtered through a 100 micron cell strainer. A crude nuclei suspension was prepared from the homogenate by 1-2 rounds of washing (Nuclei PURE store buffer, Sigma, S9183) and centrifugation (700g for 3 minutes at 4 degrees). Nuclei were then labelled with a nucleic acid binding dye (DAPI, 1:500), washed once more with nuclei isolation buffer, and pelleted by centrifugation (700g, 2 minutes at 4 degrees). Final nuclei suspensions were prepared by resuspending crude pellets in nuclei isolation buffer and sorting DAPI+ events into a collection buffer (0.5% UltraPure BSA and 1x Superasin in PBS) using a BD Influx cell sorter.

### Bulk RNA library preparation and sequencing of pilot samples

A pilot study to confirm sample integrity was performed using brain samples from two individuals, Total RNAs were extracted from bulk brain tissue or nuclei (described above) using TRIzol Reagent (Invitrogen, Carlsbad, CA, USA) according to the manufacturer’s instructions. The RNA was cleaned up using Qiagen RNeasy Micro kit, appendix C, with Dnase I (Cat# 74004; QIAGEN Sciences, Germantown, MD, USA). The integrity of the and the quantification were determined using a Bioanalyzer RNA 6000 Pico chip on the Bioanalyzer 2100 (Cat# 5067-1513, Agilent Technologies, Santa Clara, CA, USA). Strand-specific RNASeq libraries were created using the NEBNext Single Cell/Low Input RNA Library Prep Kit for Illumina (NEB #6420L; New England Biolabs, Inc., Ipswich, MA, USA). The library sizes and quantification were determined using the Bioanalyzer High Sensitivity DNA kit (Cat# 5067-4626, Agilent). The libraries were sequenced using the Nextseq500 (Illumina, Inc., San Diego, CA, USA).

### Single nucleus RNAseq library preparation, data pre-processing, normalization, and cell type assignment

The isolated nuclei were prepared and subjected to snRNA-seq using Chromium single Cell 3’ Library Kit with v2 chemistry with a Chromium Controller per the manufacturer’s instructions (10x Genomics). The library was sequenced using the Nextseq500 (Illumina). Sample demultiplexing, barcode processing, and single cell counting was performed using the Cell Ranger analysis pipeline (10x Genomics). RNAseq reads were mapped to the human reference genome with pre-mRNA annotations, which account for both exons and introns. The subsequent data analysis was performed using R/Bioconductor and Partek Single Cell Toolkit (Partek). For quality control, nuclei with high mitochondrial content, high UMI, high gene number per cells were removed. Data were normalized using a scaling factor of 1,000,000. The AUCell algorithm was used to score each nuclei for enrichment of cell-type specific transcriptomes (genes described in **Supplementary Table 2**).

### Identifying of SPMS-associated microglial subtypes using graph-based clustering

The 2211 cells annotated as microglia/phagocytes were divided into four data subsets based on batch and counts. The first sequencing batch (637 cells) was filtered to cells with 800-1800 total counts, which was used as the discovery set (421 cells). The remaining 216 cells from the first sequencing batch, all 234 cells from the second sequencing batch, and all 1340 cells from the third sequencing batch were treated as separate data subsets (ie batches).For the discovery set, the count matrix was log normalized (base 2), identified the top 2000 variable genes, scaled the matrix with zero mean and 1 variance, performed Principal Component Analysis (PCA) to reduce the dimensions. Finally, Graph-based clustering was performed to derive 3 microglial clusters (g1, g2, g3). To perform clustering analysis on the entire dataset of 2211 microglia/phagocytes, R/Seurat was used. The raw gene expression matrix was log normalized using *NormalizeData* with a scale factor of 10,000. Then the top 2,000 variable genes were identified using *FindVariableGenes* and the matrix was scaled and centered to zero mean and 1 variance using *ScaleData*. PCA was performed to obtain a low-dimensional matrix. Before the graph-based clustering, *RunHarmony* with default parameters was used to remove the batch effect between four subsets. Finally, *FindNeighbors* was performed on the harmonized matrix and *FindClusters* was used to derive microglial subclusters with a resolution of 1.2. Overall, 11 microglial subclusters were identified.

### Quantification of FTL protein in tissue samples

Lysates were prepared from frozen brain homogenized in Cell Signaling lysis buffer (Cell Signaling #9803) or RIPA buffer, with protease/phosphatase inhibitors and benzonase (1:500 dilution; Sigma E1014). Samples were diluted 1:250 and loaded on the 96 well FTL ElISA plate (Abcam, Ferritin ELISA Kit, ab108698) and run as per manufacturer instructions.

### Histology and iron staining

Immunohistochemistry was performed on 30 micron brain sections using antibody against myelin proteolipid protein as previously described*. Diaminobenzidine (DAB)-enhanced Turnbull staining was performed for detection of total iron (ferric and ferrous). 30 micron brain sections were immersed in aqueous ammonium sulfide followed by incubation in aqueous potassium ferricyanide and hydrochloric acid. The staining was amplified by immersing in DAB.

### RNAscope

RNA transcripts were detected in 7 micron brain sections using the RNAscope 2.5 HD Duplex kit from Advanced Cell Diagnostics. FTL and TYROBP probes were used in channel 1 and P2RY12 probe in channel 2. Hematoxylin counterstain was used to label nuclei. Images were taken on a Zeiss Axiophot microscope with Zeiss Axiocam 512 color camera.

### Bulk RNA library preparation and sequencing of independent brain tissue sections

RNA was extracted from 25 frozen brain sections according to the “Purification of Total RNA from Microdissected Cryosections” method of the Qiagen RNeasy Micro kit, with Dnase I (Cat# 74004; Qiagen). The integrity of the and the quantification were determined using a Bioanalyzer RNA 6000 Pico chip on the Bioanalyzer 2100 (Cat# 5067-1513, Agilent). Strand-specific RNASeq libraries were created using the TruSeq stranded mRNA library prep Kit (cat# RS-122-2101, Illumina). The library sizes and quantification were determined using the Bioanalyzer High Sensitivity DNA kit (Cat# 5067-4626, Agilent). The libraries were sequenced using the HiSeq2000 (Illumina).

### Bulk RNA data processing, statistical analysis, and pathway enrichment analysis of patient tissue samples

All data processing from the QC of FASTQ files to the derivation of differentially expressed genes was performed within Array Studio (Version 10, Omicsoft Corporation, Research Triangle Park, NC, USA). Data quality was assessed using the “Raw Data QC Wizard” function within Array Studio. The sequence used to trim the adapters from the reads was “AGATCGGAAGAGCG.” Paired reads were mapped to the human reference genome (GRCh38, GenCode.V24) using the Omicsoft Aligner 4 (OSA4)^29^. The EM algorithm was implemented to calculate the FPKM (Fragments Per Kilobase Million) value for each gene^30^. Lowly expressed genes were filtered out. Additionally, the genes were filtered to retain only protein coding genes. Next, the samples were normalized (upper quantile) followed by a log2 transformation (Log2[FPKM+1]). For the independent set of brain samples, the Students T-tests was performed to determine which genes were significantly differentially expressed between groups. Genes were considered significantly altered if p-value <0.05 and the absolute fold-change in expression was ≥1.2.Differentially expressed genes (Log2 fold change cut off of 0.25, adjusted p-value <0.05) were used to determine canonical pathway enrichment (Ingenuity Pathway Analysis, Qiagen).

### Human monocyte isolation, differentiation, and culture

Human peripheral blood mononuclear cells (PBMCs) were isolated from Leukopaks (Fisher Hemacare Corp) using histopaque-1077 (Sigma) according to manufacturer’s protocols. Monocytes were then isolated with CD14 microbeads and the AutoMACS Pro Separator (Miltenyi Biotec) according to manufacturer’s instructions. Monocytes were plated on low cell binding culture plates at 1.2 million cells per mL in RPMI supplemented with 10% FBS and 100 ng/mL CSF-1, with half the medium replaced after 3 days. After 7 days of culture, differentiated macrophages were collected and used for experiments or cryopreserved for later use. Acute inflammatory-type macrophages were induced by treating cells with 10 ng/mL lipopolysaccharide (from E. coli 055:B5, Sigma) for 3.5 hours. Cells were lysed with RLT buffer (Qiagen) and frozen at −80 for downstream RNA isolation, library construction and sequencing (Performed by BGI Americas Corporation).

### Iron loading, viability, and lipid peroxidation experiments with human microglial cell line

The human microglia cell line (abm T3451) was plated at 5,000 cells/well in a 96-well PDL coated black optical imaging plates (Greiner #655946) plate in 100uL of complete microglia media (High glucose DMEM (Fisher scientific 10313-021) + 1x L-Glutamine (Millipore TMS-002-C) + 1x Penn Strep (Millipore TMS-AB2-C), and 10% FBS (Takarabio 631101). Cells were left cells overnight in 37C incubator. The following day, the cells were treated for 24hrs with appropriate treatment. FeSO4 (Sigma F8633-250G) was dissolved in cell culture-grade H2O (Invitrogen 10977-015) to make an 80mM solution. The 80mM iron solution was diluted 1:100 in microglia media to reach a working concentration of 800uM. It was Serial diluted 2-fold to make 400uM, 200uM, and 100uM media/iron treatments. RSL3 (Sigma SML2234) was added at 1uM. Liproxstatin-1 (Sigma SML1414) was added at 100nM to appropriate wells. Media was full exchanged to treatment media. Twenty-four hours after treatment. Cells were first analyzed for lipid peroxidation with the ImageIT kit (life technologies C10445). Image-IT dye was diluted in microglia media at 500x. 1 drop of DAPI solution (Invitrogen #R37605) was added per mL of media. 100 uL of dye+DAPI media was added to the existing 100uL in each well. Cells were incubated for 60 minutes in 37C incubator. Cells were then washed 3x in RT PBS (Gibco #20012-027) and placed in 100uL PBS. Cells were imaged on the InCell analyzer and analyzed on Cell Profiler. Immediately after imaging cell viability was measured by Cell Titer Glo (Promega #G9241). 100uL of Cell titer glo solution was added to each well. The plate was shaken on orbital shaker for 2 min at RT. The plates were then left at room temperature for 10 minutes. After ten minutes, 100uL was pulled from each well and placed into an opaque plate. Luminescence was measured on a Flexstation.

For transcriptomic studies, cells were plated at 5 million cells/ T75 flask in 13mL complete microglia media (High glucose DMEM (Fisher scientific 10313-021)+ 1x L-Glutamine (Millipore TMS-002-C)+ 1x Penn Strep (Millipore TMS-AB2-C), and 10% FBS (Takarabio 631101). Cells were left cells overnight in 37C incubator. 10x solutions were made for each condition. For vehicle, DMSO was added at 1:100 to complete microglia media for 10x solution. For iron only, FeSO4 (Sigma F8633-250G) was dissolved in cell culture-grade H2O (Invitrogen 10977-015) to make an 80mM solution. The 80mM solution was diluted 1:5 in microglia media to reach a 10x concentration of 16mM. For RSL3 only, 10mM stock RSL3 (Sigma SML2234) was added at 1:1000 to complete microglia media for a 10x solution (10uM RSL3). For Liproxstatin-1 only, 1mM stock liproxstatin-1 (Sigma SML1414) was added 1:1000 to complete microglia media for a 1uM 10x solution. For iron + RSL3, 10mM RSL3 was added 1:1000 to 16mM Fe in microglia media for a 10uM RSL3 + 16mM Fe 10x solution. For iron + RSL3 + lip-1, the iron + RSL3 solution 10x solution as mad and 1mM lip-1 was added 1:1000 for a 10uM RSL3 + 16mM Fe + 1uM lip-1 10x solution. 1.3mL were removed from each flask and 1.3mL of appropriate 10x solution was added. Cell pellets were collected at 4 hours post exposure. Media was aspirated and flasks were washed 2x with RT PBS (Gibco #20012-027). Cells were lifted with 0.25% Trypsin-EDTA (Sigma #T4049) for 5 min at 37C. An equal volume of microglia media was added to quench. Cell suspensions were spun at 1200rpm for 5min at 4C. Supernatant was aspirated, and the pellet was resuspended in PBS. Cells were spun down at 300g for 5min at 4C. Supernatant was aspirated the cell pellet was flash frozen on dry ice. Frozen pellets were stored at −80C until sent to BGI for RNAseq.

### Differential gene expression and GO analysis

The top 15 DEG from in vitro iron loading experiments were used to define a gene set reflecting iron loading with RSL3 (FE+RSL3) or vehicle alone (FE alone). Single nuclei were then scored for enrichment of these gene sets using AUCell in Partek Flow (minimum gene set of 7 of 15 in the top 10% of expressed genes).

---

**Supplemental Figure 1.**
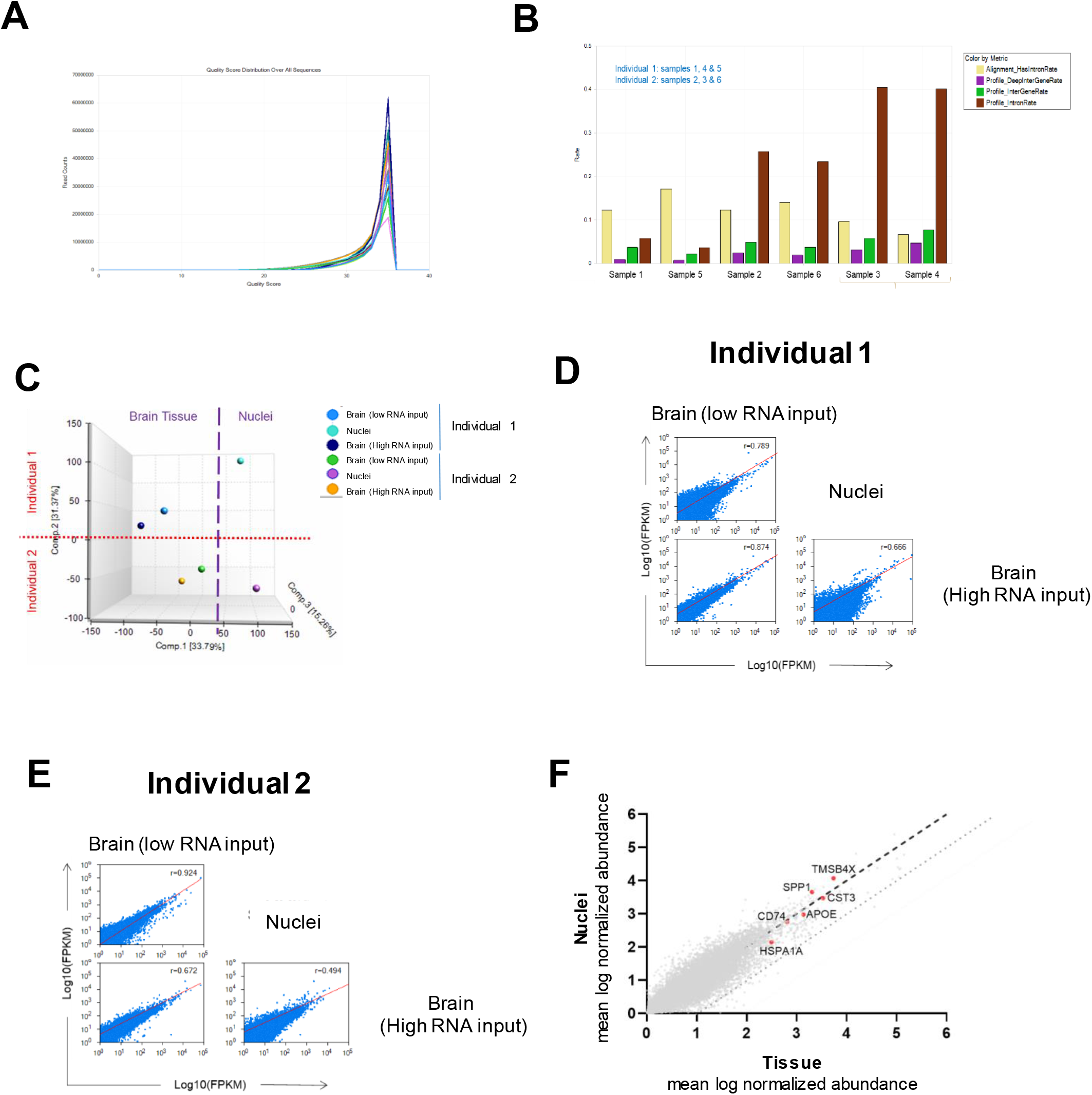
QC of FASTQ files indicates that sample quality is maintained during nuclei extraction and FACS. (**A**) Intronic & Intergenic representation across samples. (**B**) Graph of quality score distribution across all sequences indicates all samples peak at a score of 35. (**C**) PCA separates samples by input material and individual. (**D, E**) Correlation analysis demonstrates that transcript profiles are preserved between bulk tissue and bulk nuclei RNA sequencing. (**F**) Scatterplot of log normalized abundance in tissue (x-axis) versus bulk nuclei (y-axis).

**Supplemental Figure 2.**
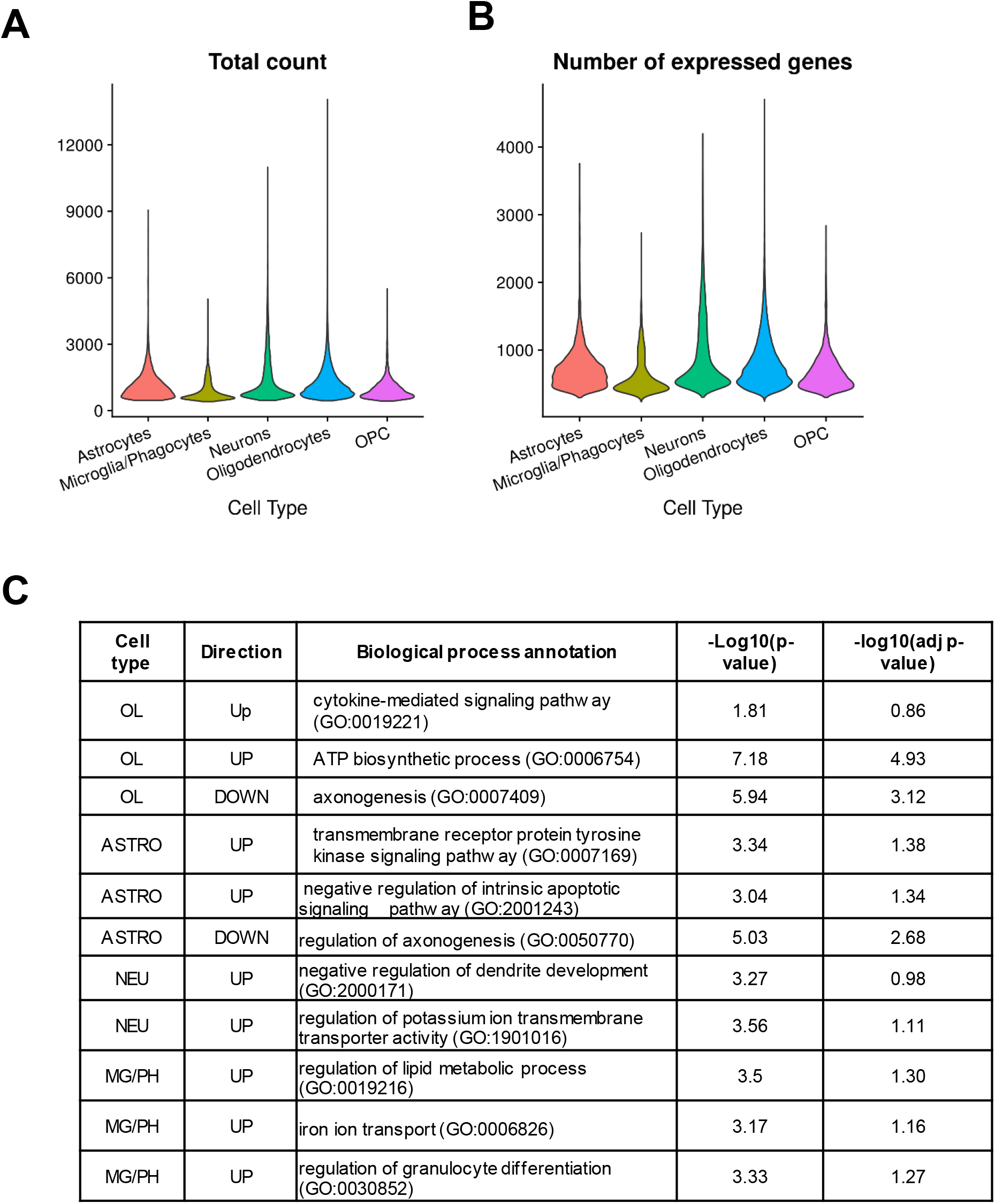
Overall detected genes by cell types. (**A**) Number of expressed genes across cell types. (**B**) Distribution of total counts across cell types. (**C**) Enriched GO BP terms among DEG by cell type. OL, oligodendrocytes; ASTRO, astrocytes; NEU, neurons; MG/PH, microglia/phagocytes

**Supplemental Figure 3.**
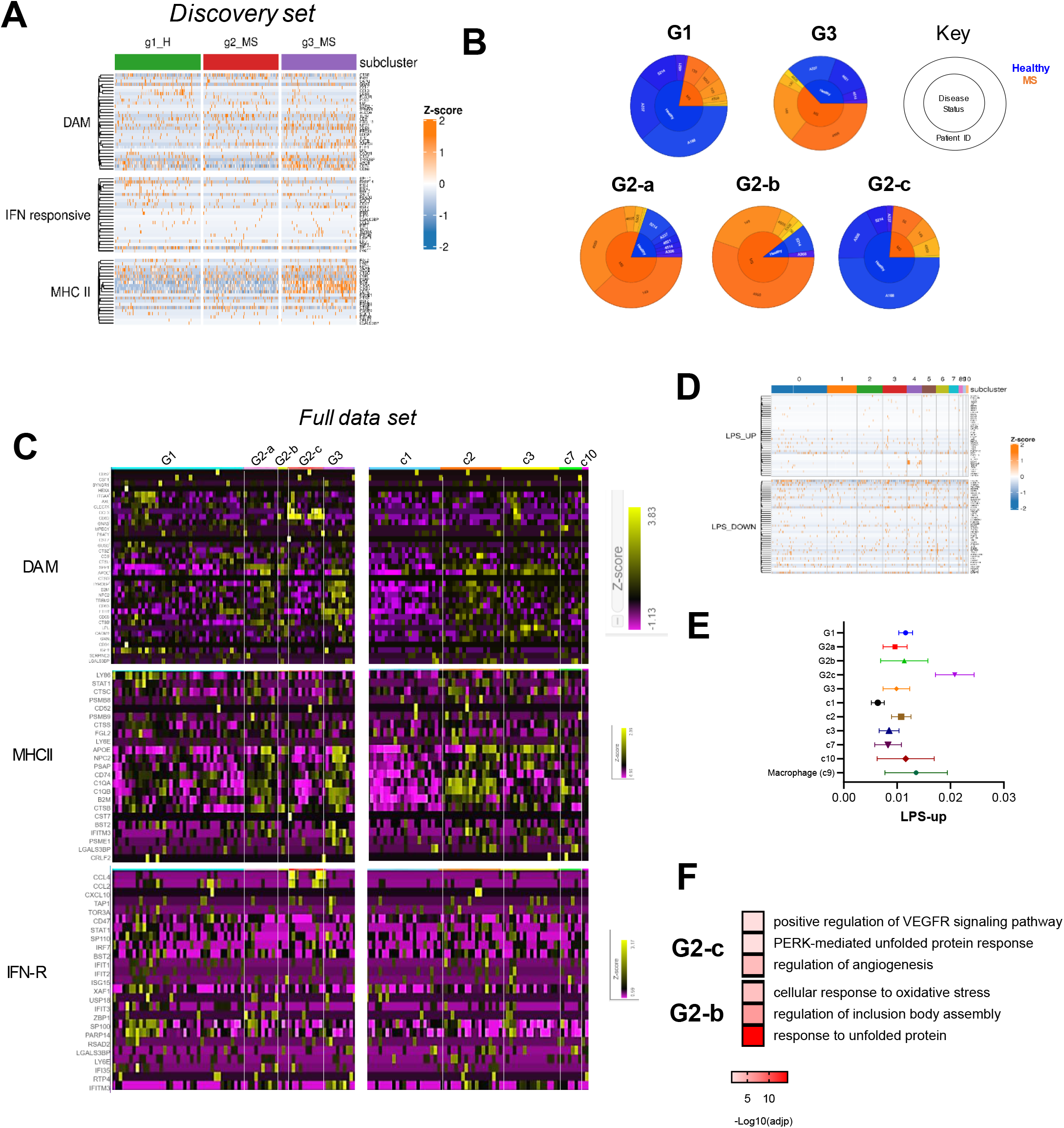
(**A**) Heat map for genes associated with murine DAM, IFN-R, and MHCII+ phenotypes in discovery set. (**B**) Pie charts of G1 (cluster 0), G2-a,b,c (clusters 4, 5, 8) and G3 (cluster 6) cell state distribution by sample (outer ring) and disease status (inner ring). (**C**) Heat map for genes associated with murine DAM, IFN-R, and MHCII+ phenotypes in all microglia clustes. (D) LPS-induced changes across clusters. **(D)** LPS-induced changes across clusters. (**E**) AUCell scoring for enrichment of LPS-up gene set (represented as mean and 95% confidence interval). (**F**) Select gene ontology annotations for DEG from G2-b and G2-c clusters. DAM, disease-associated microglia; IFN-R, interferon-responsive microglia; MHCII+, antigen presenting-like microglia.

**Supplemental Figure 4.**
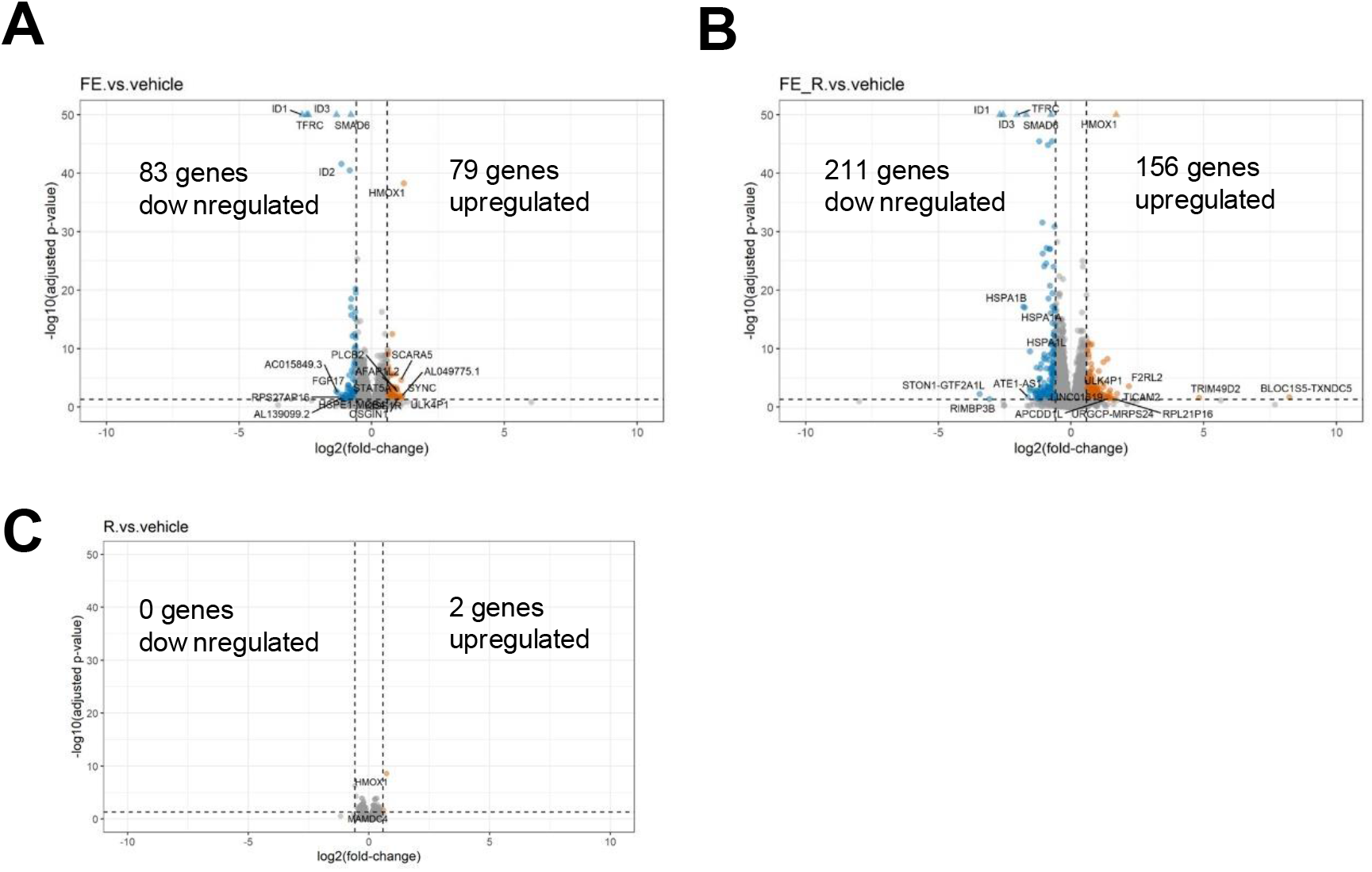
Volcano plot of DEG for cells exposed to (**A**) iron without and (**B**) with GPX4 inhibition (+RSL3). (**C**) DEG from RSL3-treated cells in the absence of iron loading. (**B**) Heat map of G1, G2a, G3, and G2/G3 shared genes across all clusters.

**Supplemental Figure 5.**
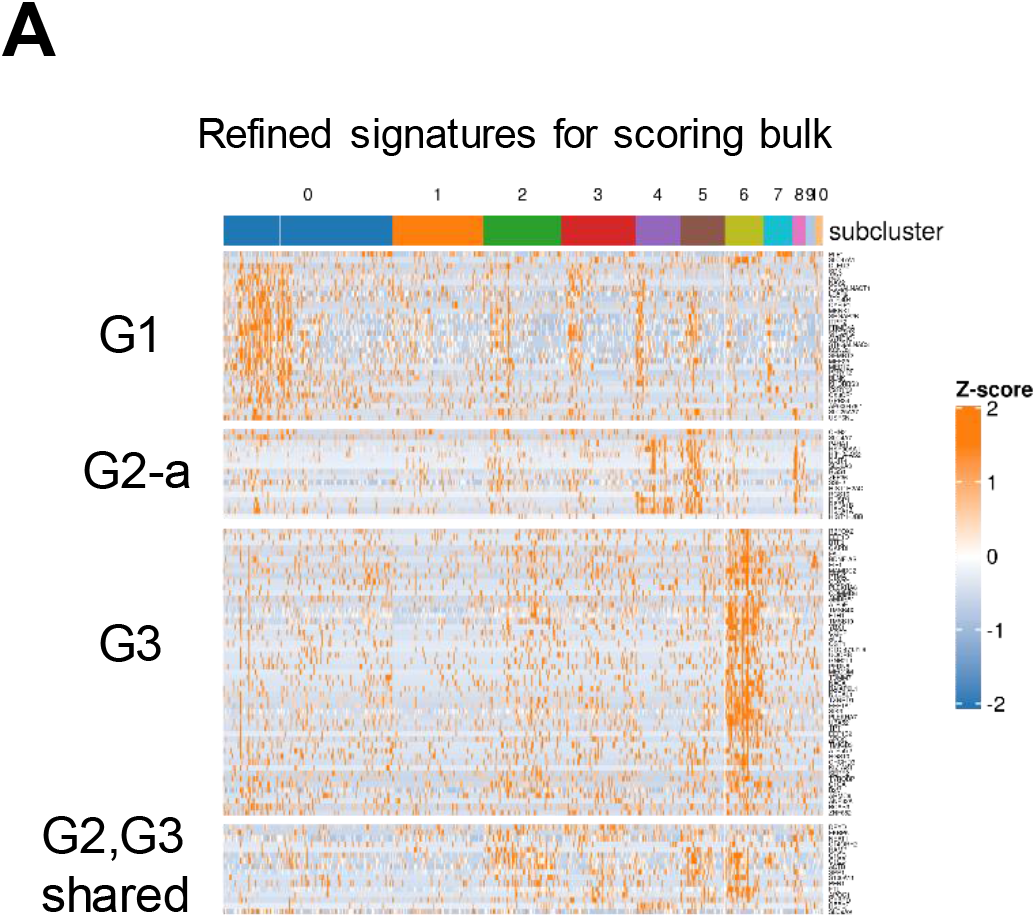
Heat map of G1, G2a, G3, and G2/G3 shared genes across all clusters.

