## Supplemental Tables for "Disrupted microglial iron homeostasis in progressive multiple sclerosis"

| Status | Patient ID | Age (yrs) | Sex | PMI (hours) |
|---|---|---|---|---|
| CTL | A306 | 65 | F | 7 |
| CTL | A337 | 86 | F | 29 |
| CTL | A368 | 65 | M | 11 |
| CTL | 4514 | 66 | M | 17.3 |
| CTL | A168 | 69 | F | 19 |
| CTL | 4621 | 83 | F | 17.6 |
| CTL | 5214 | 61 | M | 19.5 |
| CTL | 4823 | 35 | F | 11.3 |
| SPMS | 4605 | 62 | F | 11.8 |
| SPMS | 5308 | 54 | F | 9.3 |
| SPMS | 5053 | 59 | M | 24.9 |
| SPMS | 56 | 59 | F | 5.08 |
| SPMS | 130 | 40 | F | 5.25 |
| SPMS | 149 | 61 | F | 3 |
| SPMS | 4959 | 47 | F | UNK |

**Table 1.** Age, sex, and post-mortem interval (PMI) for samples used for sNuc-seq. CTL, control; SPMS, secondary progressive MS; F, female; M, male; UNK, unknown/data missing.

| **Astrocyte (21 genes)** | **Microglia (22 genes)** | **Neuron (19 genes)** | **Oligodendrocyte (20 genes)** | **OPC (13 genes)** |
|---|---|---|---|---|
| ADGRV1 | ADAM28 | CCK | APLP1 | BRINP3 |
| ANGPTL4 | APBB1IP | CDH9 | CLDN11 | DSCAM |
| APOE | ARHGAP24 | CHRM3 | CNP | LHFPL3 |
| AQP4 | C10ORF11 | CNTN5 | CTNNA3 | LRRC4C |
| CHI3L1 | C1QB | CNTNAP2 | ENPP2 | LUZP2 |
| CLU | C1QC | GABRB2 | LINC01608 | MEGF11 |
| COL5A3 | CD74 | GALNTL6 | LURAP1L-AS1 | MMP16 |
| CST3 | DOCK8 | GRIK2 | MAN2A1 | NOVA1 |
| DPP10 | FYB | GRIN2A | MOBP | PTPRZ1 |
| DTNA | INPP5D | KCNC2 | MOG | RP4-668E10.4 |
| GFAP | LPAR6 | KCNQ5 | PIP4K2A | SEMA5A |
| GLIS3 | MEF2A | MEG3 | PLCL1 | TNR |
| GPC5 | P2RY12 | RBFOX1 | RNF220 | VCAN |
| GPM6A | PLXDC2 | ROBO2 | SEPP1 | |
| MGST1 | PTPRC | RP11-123O10.4 | SLC44A1 | |
| RFX4 | RGS1 | SNAP25 | SLC5A11 | |
| RNF219-AS1 | RP11-624C23.1 | SYNPR | ST18 | |
| SLC14A1 | SFMBT2 | TENM2 | TMEM144 | |
| SLC1A2 | SRGN | ZNF385D | TMTC2 | |
| SPARCL1 | ST6GAL1 | | TTLL7 | |
| TPD52L1 | TBXAS1 | | | |

**Table 2.** Core gene signatures for used for annotation of astrocytes, microglia, neurons, oligodendrocytes, and oligodendrocyte progenitor cells (OPC).

| Status | Brain ID | Sex |
|---|---|---|
| CTL | 4307 | M |
| CTL | 4823 | F |
| CTL | 5214 | M |
| CTL | 4621 | F |
| CTL | 4514 | M |
| CTL | A168 8-1 | F |
| CTL | A0212 12-4 | M |
| SPMS | 5252/Sec8 | F |
| SPMS | 5053 | M |
| SPMS | 4605 | F |
| SPMS | 4399/Sec5 | M |
| SPMS | MS137 4-IRT | F |
| SPMS | 4732/Sec7 | F |
| SPMS | 5276 | F |
| PPMS | 4961 | F |
| PPMS | 4951 | F |
| PPMS | 2485 | M |
| PPMS | 5149 | M |
| PPMS | 3816 | F |
| RRMS | 5095 | M |
| RRMS | 505 | F |
| RRMS | 5154 | M |
| RRMS | 5139 | M |
| RRMS | 5102 | F |
| RRMS | 5123 | F |

**Table 3.** Samples used for bulk RNA-sequencing. CTL, control; SPMS, secondary progressive MS; PPMS, primary progressive MS; RRMS, relapsing-remitting MS.

| ID | Platform | Case.Disease |
|---|---|---|
| GSE36980.GPL6244.test3 | GPL6244 | Alzheimer's disease (AD) |
| GSE48350.GPL570.test2 | GPL570 | Alzheimer's disease (AD) |
| GSE48350.GPL570.test3 | GPL570 | Alzheimer's disease (AD) |
| GSE8397.GPL96.test1 | GPL96 | Parkinson's disease (PD) |
| GSE36980.GPL6244.test2 | GPL6244 | Alzheimer's disease (AD) |
| GSE48350.GPL570.test4 | GPL570 | Alzheimer's disease (AD) |
| GSE13162.GPL571.test2 | GPL571 | frontotemporal lobar degeneration (FTLD) |
| GSE84422.GPL570.test2 | GPL570 | Alzheimer's disease (AD) |
| E-MEXP-2280.A-AFFY-44.test2 | AFFY-44 | frontotemporal lobar degeneration (FTLD) |
| GSE84422.GPL570.test1 | GPL570 | Alzheimer's disease (AD) |
| GSE13162.GPL571.test3 | GPL571 | frontotemporal lobar degeneration (FTLD) |
| GSE39420.GPL11532.test1 | GPL11532 | Alzheimer's disease (AD) |
| GSE28146.GPL570.test1 | GPL570 | Alzheimer's disease (AD) |
| GSE7621.GPL570.test1 | GPL570 | Parkinson's disease (PD) |
| GSE84422.GPL96.test9 | GPL96 | Alzheimer's disease (AD) |
| GSE20164.GPL96.test1 | GPL96 | Parkinson's disease (PD) |
| E-MEXP-2280.A-AFFY-44.test4 | AFFY-44 | frontotemporal lobar degeneration (FTLD) |
| GSE13162.GPL571.test5 | GPL571 | frontotemporal lobar degeneration (FTLD) |
| GSE84422.GPL96.test8 | GPL96 | Alzheimer's disease (AD) |
| GSE49036.GPL570.test3 | GPL570 | Parkinson's disease (PD) |
| GSE84422.GPL96.test17 | GPL96 | Alzheimer's disease (AD) |
| GSE28146.GPL570.test3 | GPL570 | Alzheimer's disease (AD) |
| E-MEXP-2280.A-AFFY-44.test1 | AFFY-44 | Alzheimer's disease (AD) |
| GSE54562.GPL6947.test1 | GPL6947 | major depressive disorder |
| GSE49036.GPL570.test2 | GPL570 | Parkinson's disease (PD) |
| GSE1297.GPL96.test1 | GPL96 | Alzheimer's disease (AD) |
| GSE39420.GPL11532.test2 | GPL11532 | Alzheimer's disease (AD) |
| GSE8397.GPL96.test3 | GPL96 | Parkinson's disease (PD) |
| GSE29378.GPL6947.test2 | GPL6947 | Alzheimer's disease (AD) |
| GSE5389.GPL96.test3 | GPL96 | bipolar disorder |
| GSE13162.GPL571.test1 | GPL571 | frontotemporal lobar degeneration (FTLD) |
| GSE44593.GPL570.test1 | GPL570 | major depressive disorder |
| GSE28146.GPL570.test2 | GPL570 | Alzheimer's disease (AD) |
| GSE36980.GPL6244.test1 | GPL6244 | Alzheimer's disease (AD) |
| GSE54566.GPL570.test1 | GPL570 | major depressive disorder |

**Table 4.** Studies used for GSVA analysis.
